# Zebrafish cornea formation and homeostasis reflect robustness in camera-type eye biology, regardless of its environment

**DOI:** 10.1101/2022.03.28.486026

**Authors:** Kaisa Ikkala, Sini Raatikainen, Frederic Michon

## Abstract

The cornea is the transparent tissue covering the camera-type eyes. This position makes this tissue prone to environmental aggressions, and physical alteration can have a drastic negative impact on sight. The current knowledge on zebrafish corneal physiology is too scarce to get an understanding on its strategy to maintain a clear sight in aquatic environment. In this study, we focus on corneal formation and maturation in zebrafish. After describing the morphological changes taking place from 3 days post fertilization to adulthood, we analyzed cell proliferation. We show that label retaining cells appear just before 1 month of age. Our cell proliferation study, combined to the study of *Pax6a* and *krtt1c19e* expression, demonstrate a long maturation process. This process ends with a solid patterning of corneal innervation. Finally, corneal abrasion wound healing process recapitulates the maturation process, via a plasticity period. Altogether, our study decipher the maturation steps of an aquatic cornea. These findings will facilitate the use of zebrafish as model of corneal physiology studies.

## Introduction

The cornea is the transparent surface of the eye. Like skin, it serves as a barrier, protecting the eye from the external environment. In addition, the cornea is an important refractive structure of the light path, and has therefore to remain transparent to preserve sight. To retain transparency, the corneal epithelium renews constantly. Locally, from the progenitor cells on the basal layer of the epithelium (Dhouailly et al., 2014), and globally from the stem cells in the limbus, located between cornea and conjunctiva (Schermer et al., 1986; Cotsarelis et al., 1989; Lehrer et al., 1998).Due its easy access, the cornea provides an excellent model to study epithelial cell dynamics in homeostasis and upon injury.

Because of its conserved function, the corneal simple structure forms similarly across camera-type-eyed animals. The corneal epithelium develops from the surface ectoderm of the embryo. Corneal epithelial progenitor cells are located above the lens after its invagination (Dhouailly et al., 2014). After lens detachment from the ectoderm, mesenchymal cells of neural crest origin migrate below the presumptive corneal epithelium and give rise to the corneal endothelial cell layer and stromal keratocytes (Meyer and O’Rahilly, 1959; Pei and Rhodin, 1970; Johnston et al., 1979). As the cornea matures, the stroma thickens and the epithelium stratifies. The human corneal epithelium stratifies from two to five layers upon eyelid opening before birth (Sevel and Isaacs, 1988). In rodents, the stratification from a couple of epithelial layers to adult thickness occurs post-natally, by four weeks of age (Chung et al., 1992; Song et al., 2003), and is likewise considered to be stimulated by eyelid opening (Zieske, 2004). The fully stratified corneal epithelium consists of the basal cell layer, the overlying superficial cells, and the terminally differentiated outermost cells.

The differentiation status of epithelial cell populations is traditionally evaluated through the keratins that are expressed. The immature corneal epithelium is positive for *keratin 14* (*krt14*) (Richardson et al., 2017). The mature and terminally differentiated corneal epithelium is positive for *krt12*, co-expressed with *krt3* in human, and rabbit as well as chicken cornea (Kasper et al., 1988; Chaloin-Dufau et al., 1990). The shift in *keratin* expression, from *krt14* to *krt12*, reflects the epithelium maturation process (Dhouailly et al., 2014). *Keratin 19* expression, on the other hand, was reported in the basal cells in limbal and peripheral regions in human, and only limbal in mouse (Yoshida et al., 2006). More recently, we showed that *krt8* and *krt19* expression in murine cornea changed from global expression perinatally to limbal only at 3 weeks after birth (Kalha et al., 2018).

Maturation, growth, and homeostatic renewal in adult tissue require controlled proliferation. Studies of the corneal epithelial clonal pattern, indicated a switch from local to limbal renewal in three to five weeks old mice (Collinson et al., 2002; Endo et al., 2007; Zhang et al., 2008), suggesting the establishment and activation of the limbal stem cell niche. Analysis on cell proliferation on central versus peripheral / limbal areas in mice of 0 to 24 weeks old showed that limbus contained the highest, the periphery the second highest, and central cornea the lowest, number of Ki67-positive cells, throughout the stages studied (Kalha et al., 2018). Even in the embryonic stages E13.5 to E16.5, BrdU-labeling showed highest proliferation rate in the peripheral regions (Collinson et al., 2002). Collectively, these data suggest the future limbal region is relatively active in cell proliferation well before the transition from local to limbal renewal emerges, and the central cornea retains a minimal proliferative activity in its mature state.

Cornea is the most densely innervated non-neural tissue in the whole body. While the innervation process and final pattern differ between species (Lwigale, 2001; McKenna and Lwigale, 2011), the neurotrophic factors from the nerves are necessary to maintain the corneal integrity. Therefore, corneal innervation is one of the crucial elements sustaining corneal homeostasis and transparency. The loss of corneal innervation density leads to neutrophic keratitis, which is among the causes for corneal blindness (Jabbour et al., 2021).

Regarding the genetic network that regulates eye development, PAX6 (paired box protein 6) is one of the central transcription factors (Hill et al., 1991; Jordan et al., 1992; Quiring et al., 1994; Chow et al., 1999). In the corneal context, *Pax6* is expressed from early specification through development to adulthood (Koroma et al., 1997) and was shown to promote the expression of the cornea-specific *krt12* (Liu et al., 1999). Silencing or knockout of PAX6 in cultured corneal epithelial cells decreased the expression of *krt3* and *krt12* and increased the expression of skin-specific keratins *krt1* and *krt10* (Ouyang et al., 2014; Kitazawa et al., 2019), indicating Pax6 is necessary for the maintenance of the differentiated state in corneal epithelium. The loss of one pax6 allele, such as seen in the context of the rare disease called aniridia, as well as *Pax6* overexpression showed overlapping effects, such as reduced *krt12* expression, reduced adult corneal diameter or thickness, and altered wound healing (Ramaesh et al., 2003; Dora et al., 2008). Thus, an appropriate level of Pax6 is necessary to maintain corneal integrity, as we presented previously (Ikkala et al., 2022).

During terrestrialization (Precambrian – Devonian), the corneal environment changed drastically, as the eye surface became exposed to a drier habitat. This change is reflected in the maturation process of the corneal epithelium (Zieske, 2004). In aquatic species such as zebrafish, the development, maturation, and growth of the cornea proceeds while being immerged, and without the influence of the tear film. It is thus of great interest to compare corneal renewal in aquatic and terrestrial animals. Pan and colleagues reported a shift from patchy clonal pattern to limbal stripes in 4 weeks old zebrafish (Pan et al., 2013), suggesting similar maturation and adult renewal dynamics as seen in mice. Detailed studies on proliferation, and molecular cues on the maturation process of the zebrafish cornea are still scarce.

Here, we describe zebrafish corneal epithelium maturation and growth with respect to corneal morphology, apical cell appearance, proliferation, and innervation pattern. In addition, we show the expression patterns of *pax6a*, one of the zebrafish genes coding for the paired box protein 6 (Krauss et al., 1991), and *krtt1c19e*, a type 1 keratin previously reported to be expressed in the basal epidermis in zebrafish (Lee et al., 2014). Together, these data depict the corneal epithelial maturation process. Finally, we describe the effect of an injury on the molecular parameters.

## Results

### Corneal morphogenesis exhibits a long maturation process in zebrafish

Earlier studies using detailed analyses with transmission electron microscopy showed that by 3dpf, the zebrafish cornea contains all three main compartments: the endothelium, the stroma, and the epithelium (Soules and Link, 2005; Zhao et al., 2006). We investigated the overall morphology of zebrafish cornea during its morphogenesis, from 3dpf to the adult, with histological sections (Fig. 1), and scanning electron microscopy (SEM) (Fig. 2).

**Figure 1.**
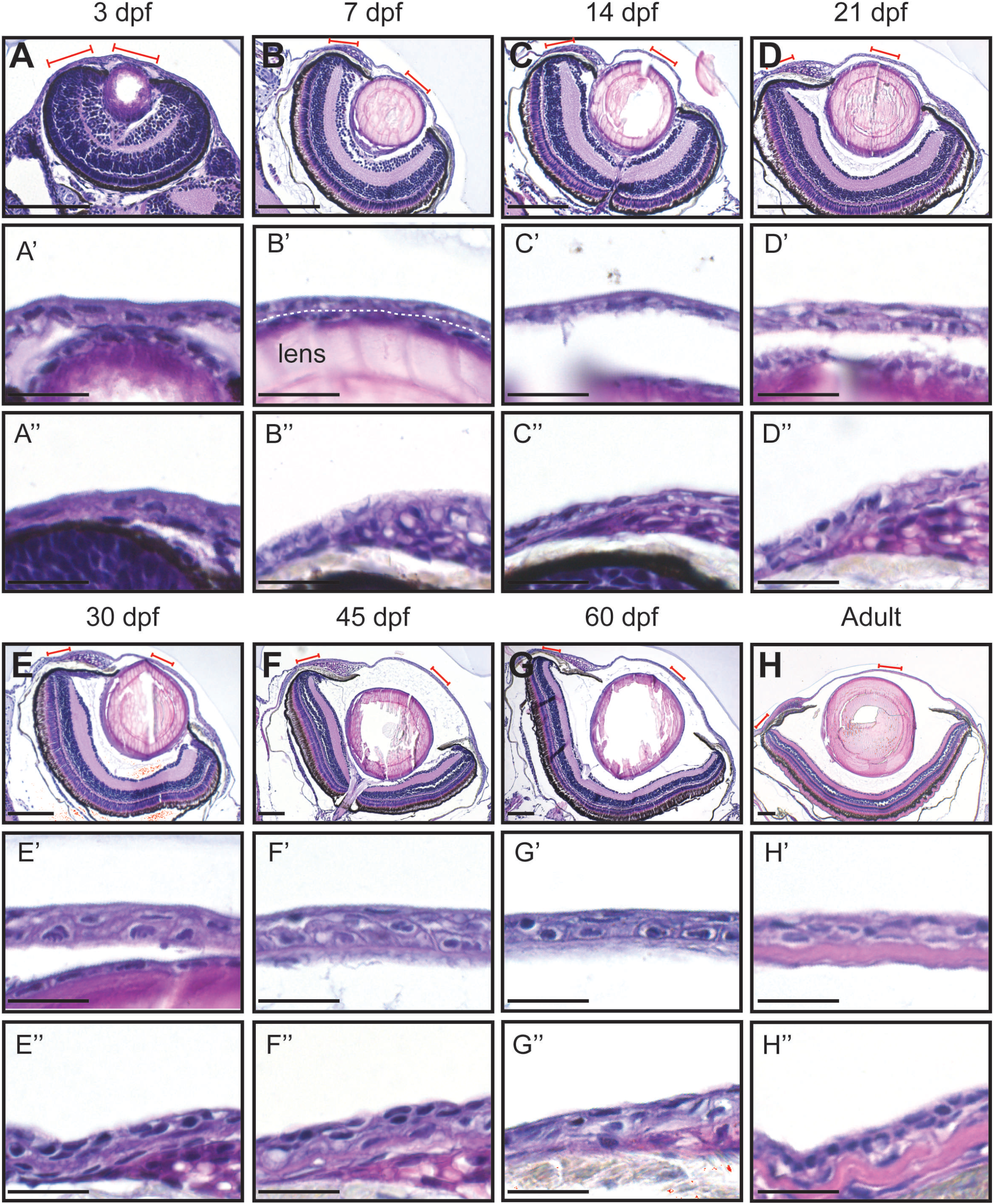
Hematoxylin-eosin staining of the eye in fish of different age (dpf, days post-fertilization). A-H: Coronal sections of the eye, anterior side on the right. Red lines indicate the regions presented in higher-magnification images. White dashed line in B’ indicates the border between the lens and the cornea. A’-H’: Cornea in the central region. A’’-H’’: Cornea in the periphery/limbus. Scale bars: 100 µm in A-H, 20 µm in A’-H’’.

**Figure 2.**
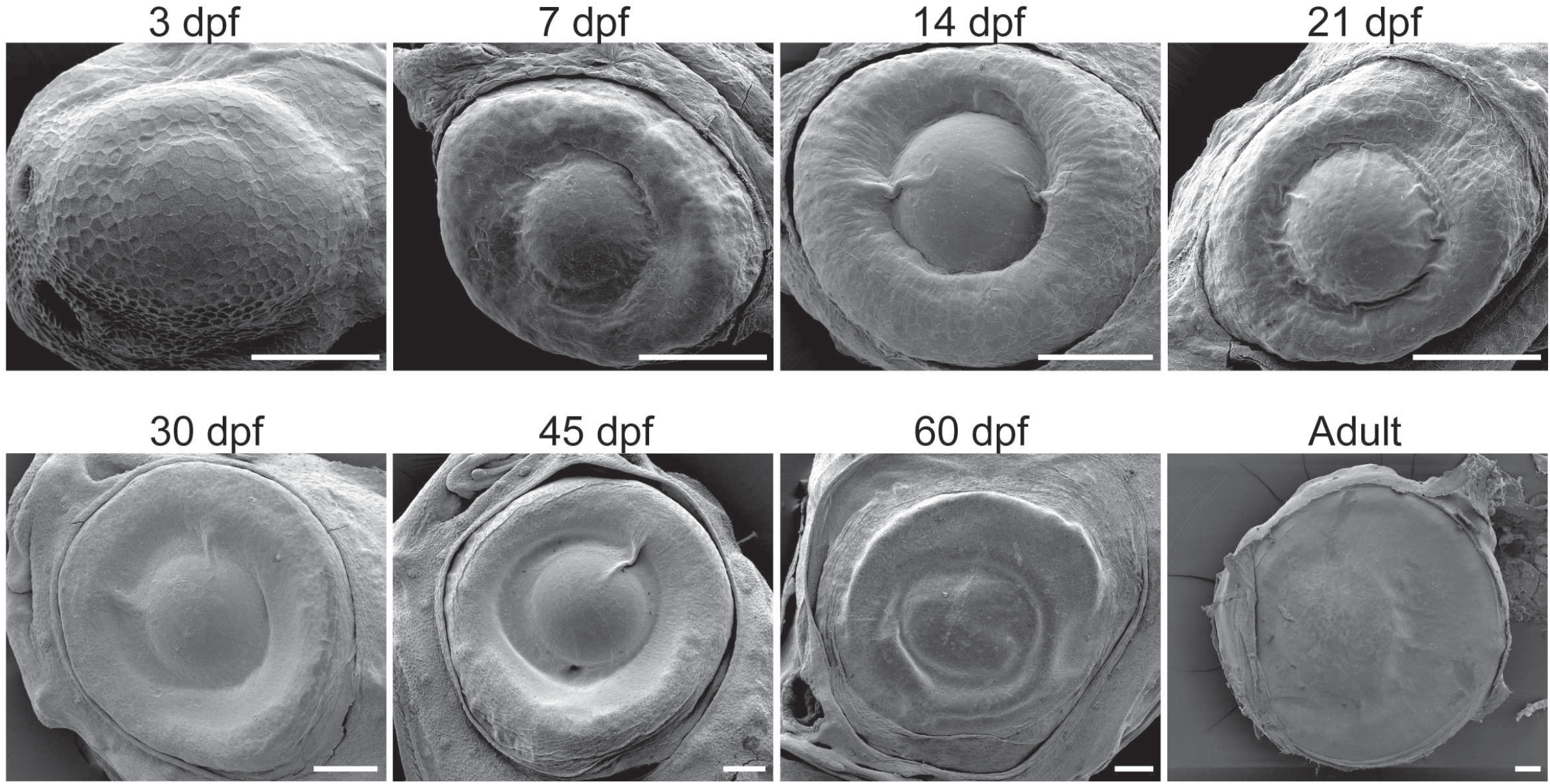
Representative SEM images of the whole eye in fish of different age (dpf, days post-fertilization). Scale bars: 100 µm.

At 3dpf, the cornea exhibited a monostratified corneal epithelium. The cornea was attached adherent to the forming lens (Figure 1A-A’). Furthermore, the peripheral area was in the continuity of the skin (Fig. 1A’’). At 7dpf, the cornea detached from the lense (Fig. 1B-B’). Furthermore, at this stage, the eye is individualized in its socket, leading to the conjunctival formation. Concomitantly, the limbus region became more complex (Fig. 1B’’). At 14dpf, the most striking changes was the increase of eye anterior chamber volume (Fig. 1 C-C’), and limbus increasing organization (Fig. 1C’’). From 21dpf to 60dpf, central and peripheral corneal domains were showing signs of maturation, such as epithelial stratification, and limbus organization (Fig. 1D-G’’). Finally, only at the adult stage the stroma was large enough to be measured (Fig. 1H-H’’).

The visualization of the eye external morphology, using SEM, confirmed some of these observations (Fig. 2). For instance, the cornea-skin continuity was lost at 7dpf. Furthermore, the lens detachment and eye anterior chamber formation might have modified drastically the eye surface morphology. Finally, the bulging of the central part of the eye was progressively loss after 60dpf, reflecting a long maturation process.

### Cell parameters confirmed the slow maturation process in zebrafish cornea

The visualization of the eye surface with SEM hinted towards a drastic eye size increase and a simultaneous cell morphology change (Fig. 2). Previously, we noticed a clear distinction between central and peripheral cell surface appearance in adult zebrafish (Ikkala et al., 2022). In the periphery, the apical cell area was remarkably smaller and the microridges more abundant compared to the central cornea. This difference could reflect the homeostatic renewal dynamics of the epithelium: on central cornea, the superficial cells are flatter and lose the microridges as they desquamate off the eye surface. Here, we studied when this difference between corneal regions occurs. We studied age groups ranging from 3 dpf to adulthood. At 3 dpf, when the eye surface was still continuous with the surrounding skin, and the superficial cells had a polygonal shape with little size variation (Fig. 3). At 7dpf, the cell-cell junctions and microridges became less striking, and the central cells began increasing in their apical area, compared to the peripheral region. During the following two weeks, the peripheral cells became heterogeneous in their microridge appearance and size. Around 21dpf, we noticed a rosette-like cell arrangement on central cornea in some samples, as reported in a previous study (Pan et al., 2013), indicating cell protrusion (Fig. S1). From 30dpf onwards, the difference between center and periphery became more obvious, the central corneal cells being larger and more heterogeneous in their size. Additionally, the peripheral cells showed abundant microridges.

**Figure 3.**
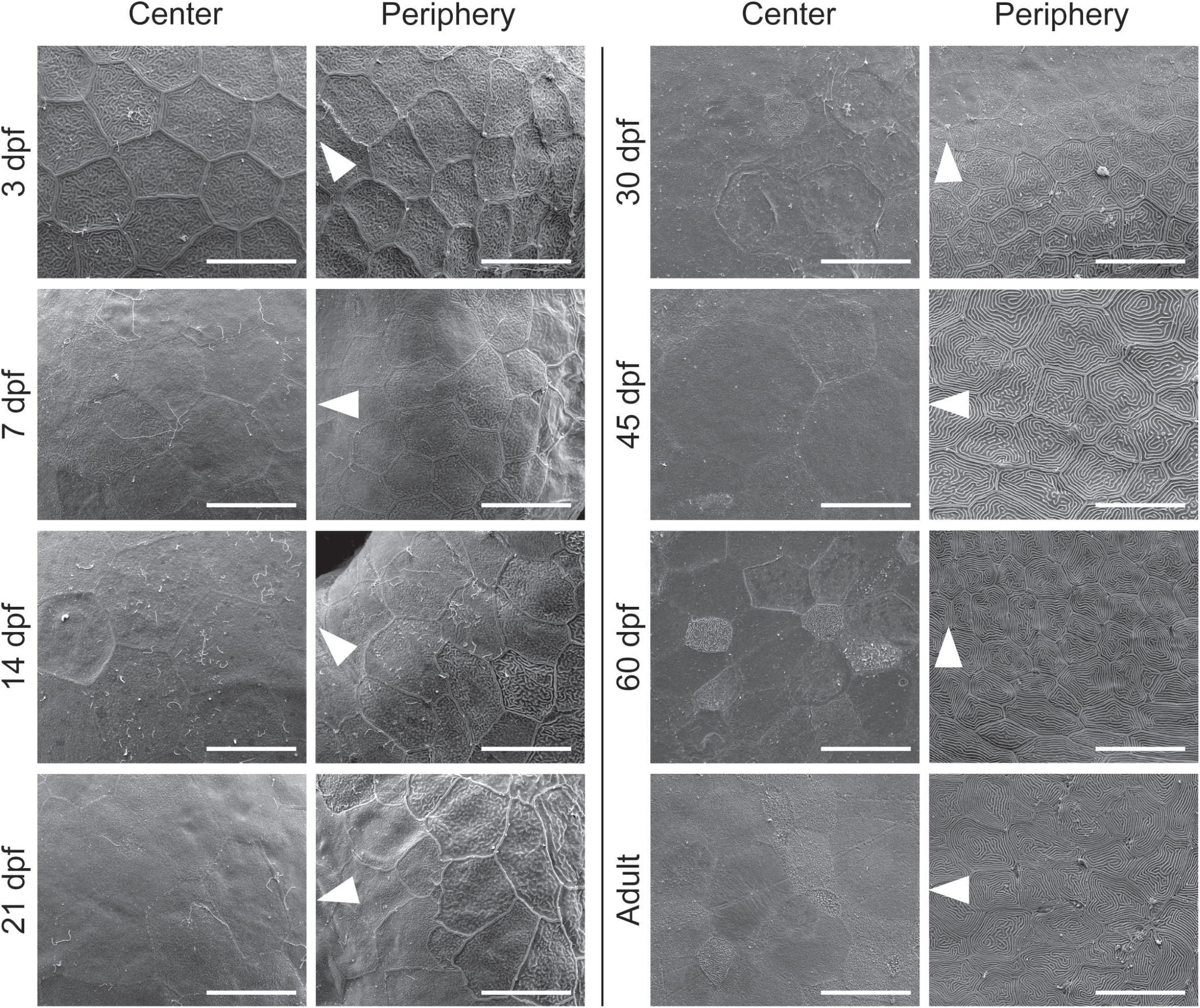
Representative SEM images of the central and peripheral regions of the eye surface in fish of different age (dpf, days post-fertilization). The white arrowheads point to the center of the eye. Scale bars: 20 µm.

To gain a better overview of the corneal growth, we quantified total fish length, eye diameter, and corneal diameter (Fig. 4). We observed modest, steady growth in total length from 3 to 30 dpf (from 3.3±0.2 to 6.3±1.3 mm), followed by more rapid growth between 30 and 45 (9.6±0.9mm) dpf, and again between 60 dpf (10.7±2.3 mm) and adult stage (23.1±1 mm) (Fig. 4A). The eye diameter followed a similar trend in growth speed (Fig. 4B), increasing from 0.4±0.0 to 0.5±0.1 mm by 30 dpf, then to 0.9±0.1 mm at 45 dpf (0.9±0.2 mm at 60 dpf), and 1.7±0.1 mm at adult stage. This would suggest that cornea is likewise going through rapid growth between 30 and 45 dpf, and still growing significantly from 60 dpf to adulthood. Since corneal total area is not solely defined by the eye diameter but also by the curvature of the corneal surface, we measured the corneal diameter along its anterior-posterior axis. When normalized to the axis of the eye, the corneal diameter increased very modestly from 3 to 60 dpf (Fig. 4C). This observation is in line with the flat shape of the zebrafish cornea, especially when compared to human or mouse cornea. Hence, corneal growth in zebrafish is likely to follow the growth speed of the eye.

**Figure 4.**
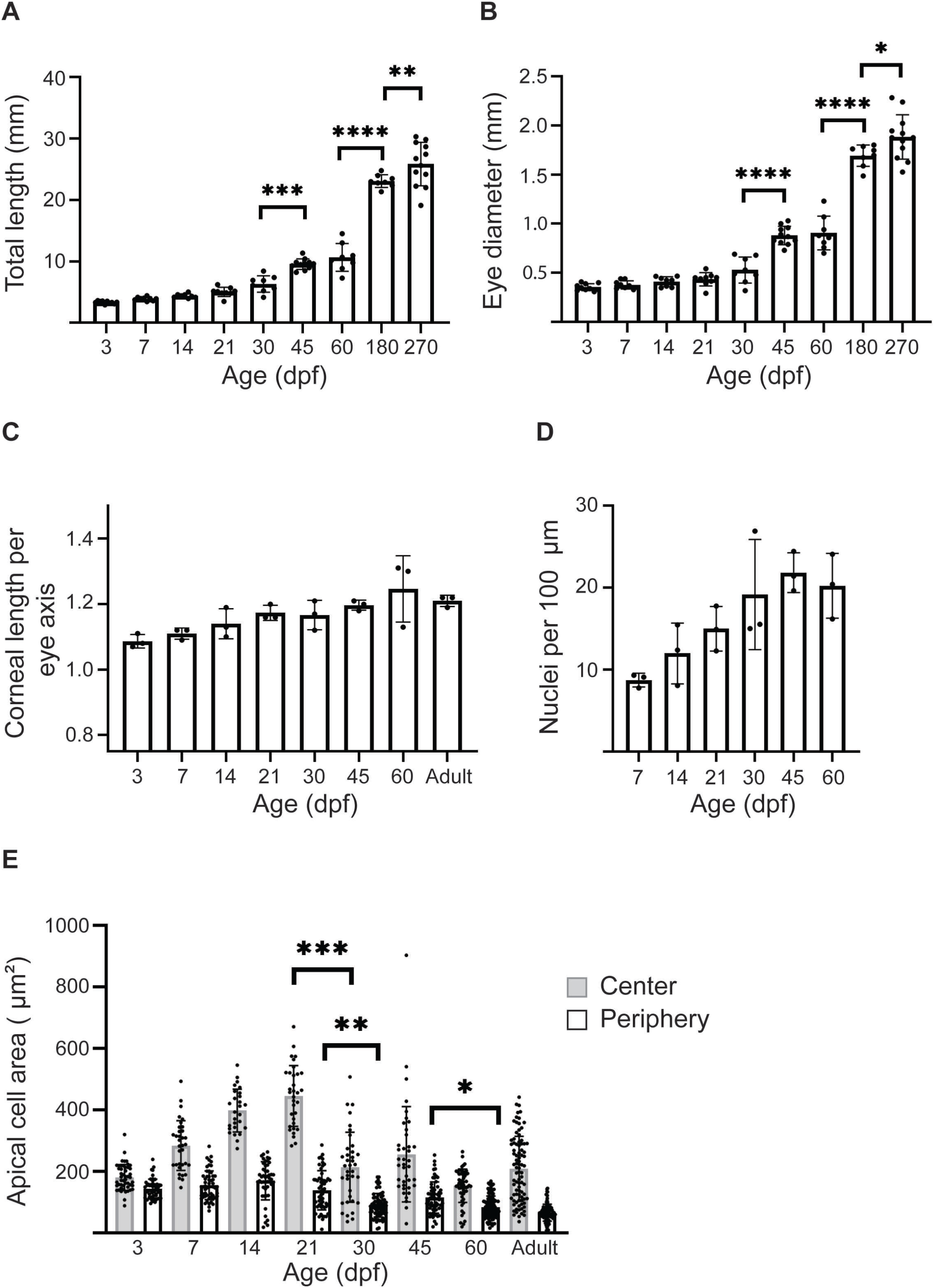
Growth of eye and corneal diameter from 3dpf to adulthood. A, B. Total length (A) and eye diameter on anterior-posterior axis (B) (n=8—12, one-way ANOVA with Sidak’s multiple comparisons test). C. Corneal length on anterior-posterior axis, normalized to anterior eye diameter (n=3, Kruskal-Wallis test with Dunn’s multiple comparisons test). D. The amount of nuclei per 100 µm on corneal epithelial section (n=3, Kruskal-Wallis test with Dunn’s multiple comparisons test). D. Apical cell area in central (gray) and peripheral (white) cornea. Cells from 3 eyes from 3 animals were pooled per age group for analysis (n=27—87 in center, 47—121 in periphery, Kruskal-Wallis test with Dunn’s multiple comparisons test). The results represent mean±stdev.

To assess the stratification process, we quantified the amount of nuclei relative to corneal diameter from 7dpf to 60 dpf (Fig. 4D). The value doubled from 0.09±0.01 at 7 dpf to 0.19±0.07 at 30 dpf (Fig. 4D), suggesting an increase in the relative amount of cells.

We also quantified the apical cell area in the SEM images (Fig. 3, 4E). Up to 21dpf, the superficial cells in the central corneal increased in average area, from 181.1±42.1 to 445.7±98.8 µm^2^, followed by a sudden decrease to 214.3±113.7 µm^2^, and remained thereafter below 260 µm^2^ (Fig. 4E). The peripheral region showed less pronounced changes, with average apical cell area above 120 µm^2^ up to 21 dpf, and below 105 µm^2^ in the following stages.

### A patterned proliferation fuels corneal growth and epithelial stratification

After observing the changes in corneal diameter and apical cell appearance, we next investigated the cell proliferation pattern with EdU and BrdU labeling during the time period between consecutive age groups from 7 to 60 dpf. EdU was administered at the beginning, BrdU at the end of each period (right before sample collection) (Fig 5A). EdU detection was not possible before 30 dpf (Fig. 5B), reflecting the dilution of EdU incorporation, meaning a high proliferative state in corneal epithelium from 3 to 21 dpf. From 30 dpf onwards, proliferating and label retaining cells were found in peripheral and central cornea, reflecting a slower proliferation starting at 21 dpf (EdU labelling for the 30dpf samples). Interestingly, the percentage of proliferating cells (Fig. 5C) increased constently from 7 (5% of BrdU+ cells) to 60 dpf (about 50% of EdU+ and BrdU+ cells). The steady increase of EdU+ cells, from 0 to 21%, reflected a change in cell fate and dynamics.

**Figure 5.**
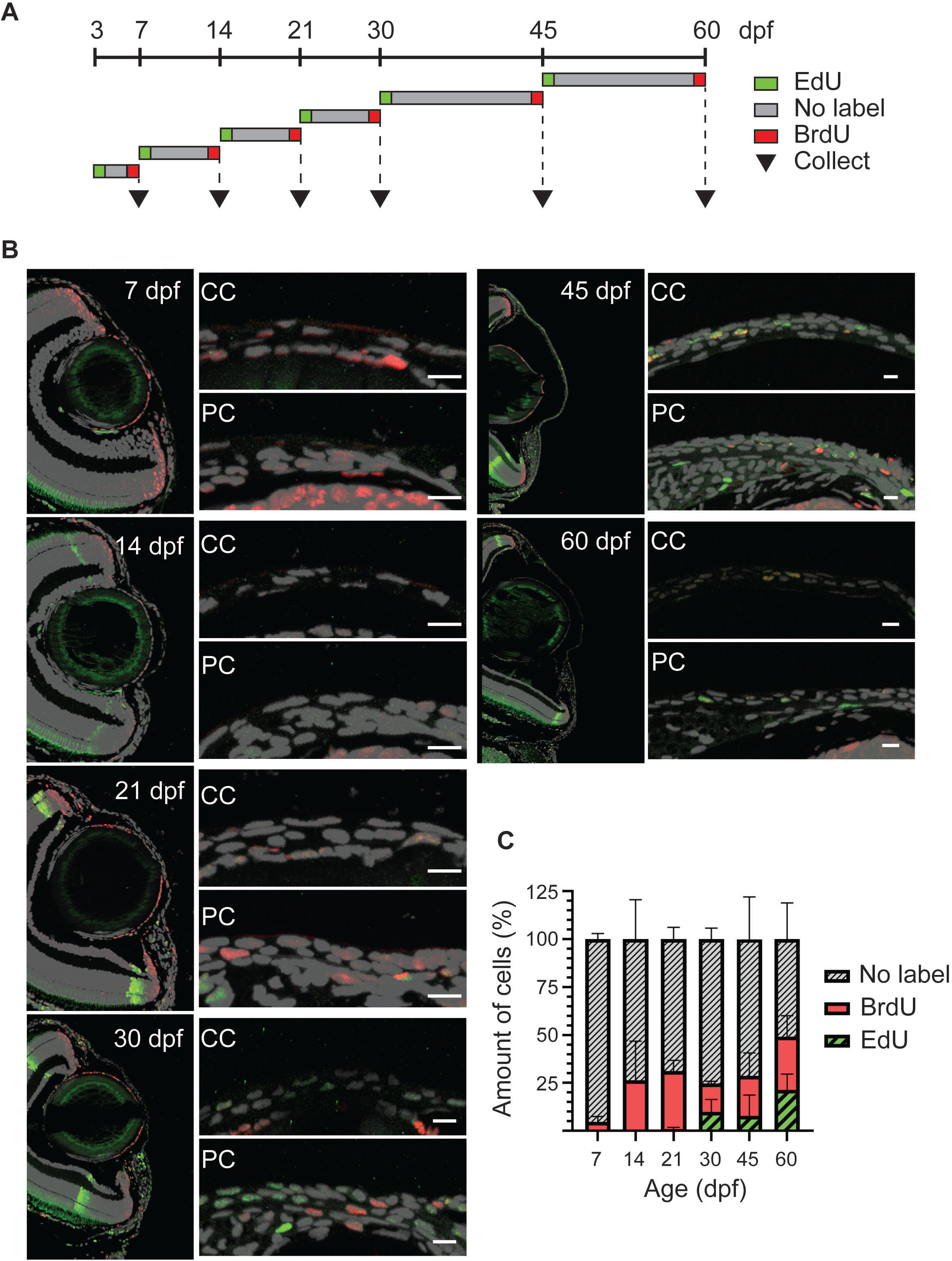
Proliferation in cornea from 3 to 60 dpf. A. A scheme showing the EdU/BrdU double labeling for each age group. B. EdU (green) and BrdU (red) staining on 5 µm paraffin sections. The panel shows the overview of the anterior eye, and central as well as peripheral/limbal regions of the cornea. C. Quantification of EdU+, BrdU+, and EdU/BrdU-cells on cornea. Results represent mean±stdev from 3 individuals per stage. The mean value of 2—3 sections from the middle of the eye were used per fish. Scale bars: 20 µm. CC= central regions, PC= peripheral/limbal region.

### Corneal innervation formation

Innervation has a critical role in corneal physiology (Yang et al., 2018). The mammalian innervation consists mostly of sensory fibers (Muller et al., 2003), which arise gradually during corneal development and maturation (McKenna and Lwigale, 2011). In murine cornea, large nerve bundles are found in the stroma, and thinner neurites emerge from the stroma to innervate the epithelium. Here, we used Acetylated Tubulin antibody staining to detect an extensive part of the nerve fiber population, as previously reported in murine cornea (Bouheraoua et al., 2019). At 3dpf, we observed neuronal processes extending on top the eye surface mainly from the posterior peripheral region (Fig. 6). Gradually, the thickest nerve trunks emerged around the peripheral cornea, giving rise to finer branches reaching towards the central region (Fig. 6-7). Notably, from 3dpf to 45dpf, the central cornea seemed weakly innervated (Fig. 6-7). Only thin fibers can be seen, and the innervation is sparce. In adult fish, there was a clear distinction between the epithelial (yellow) and stromal (red) innervation patterns (Fig. 8). Whereas the epithelial nerve branches were strongly orientated towards the center of the cornea, the stromal branches showed no specific orientation, forming a mesh. The most pronounced epithelial branches localized to the basal epithelial cell layer, indicating they would form the suprabasal nerve plexus in the zebrafish cornea. Finally, at the adult stages, we noticed thicker neurites in the central cornea, reflecting on the late maturation of corneal innervation.

**Figure 6.**
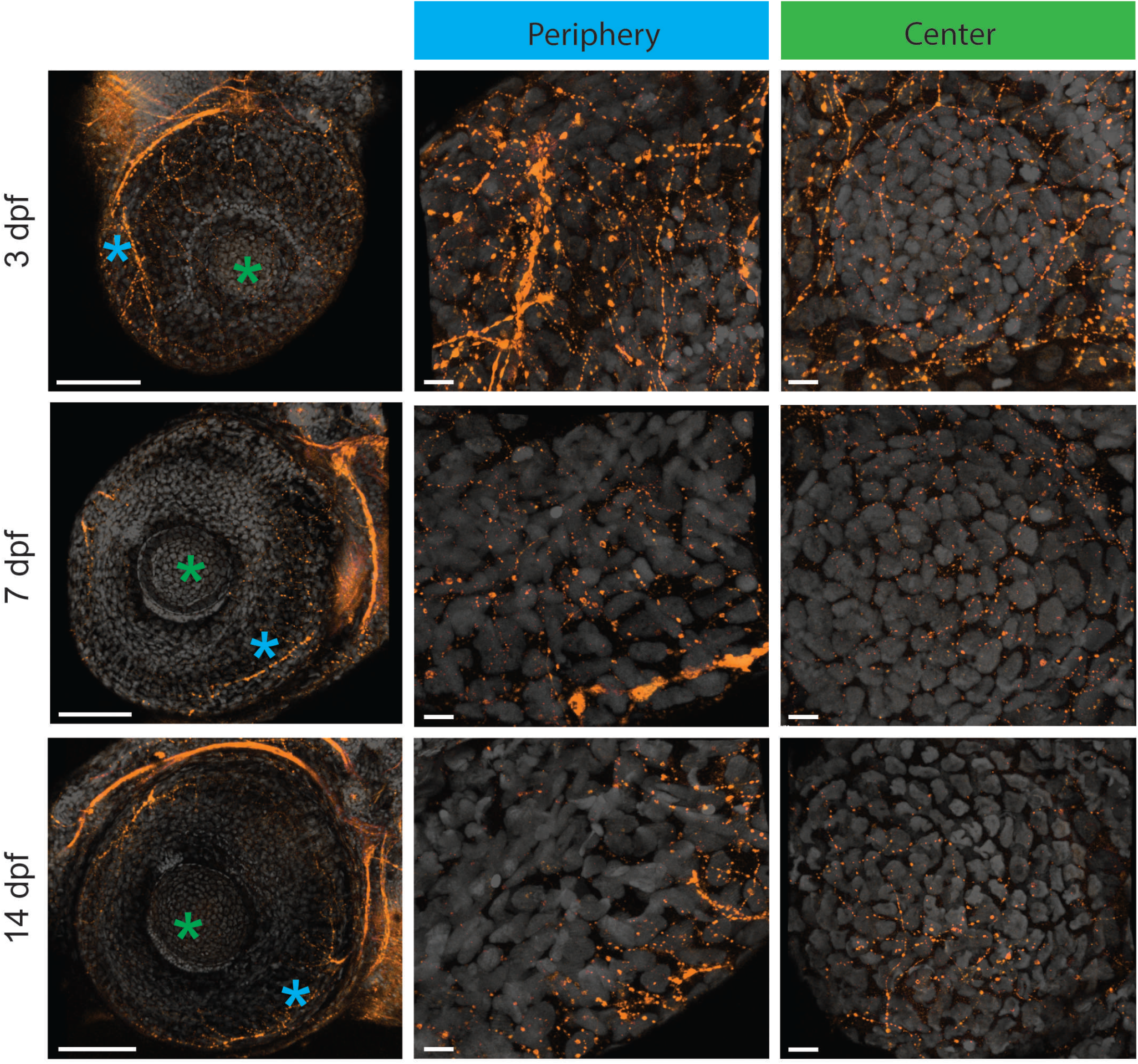
Corneal innervation before epithelial stratification at 3—14 dpf. Acetylated tubulin whole mount staining on maximum intensity projection images in central (area marked with green) and peripheral/limbal (area marked with blue) regions. During the first two weeks, the thickest neuronal branches can be observed in the peripheral regions. In central cornea, however, the signal intensity decreases during this time period. Scale bars: 100 µm in the overview images, 10 µm in the magnified images.

**Figure 7.**
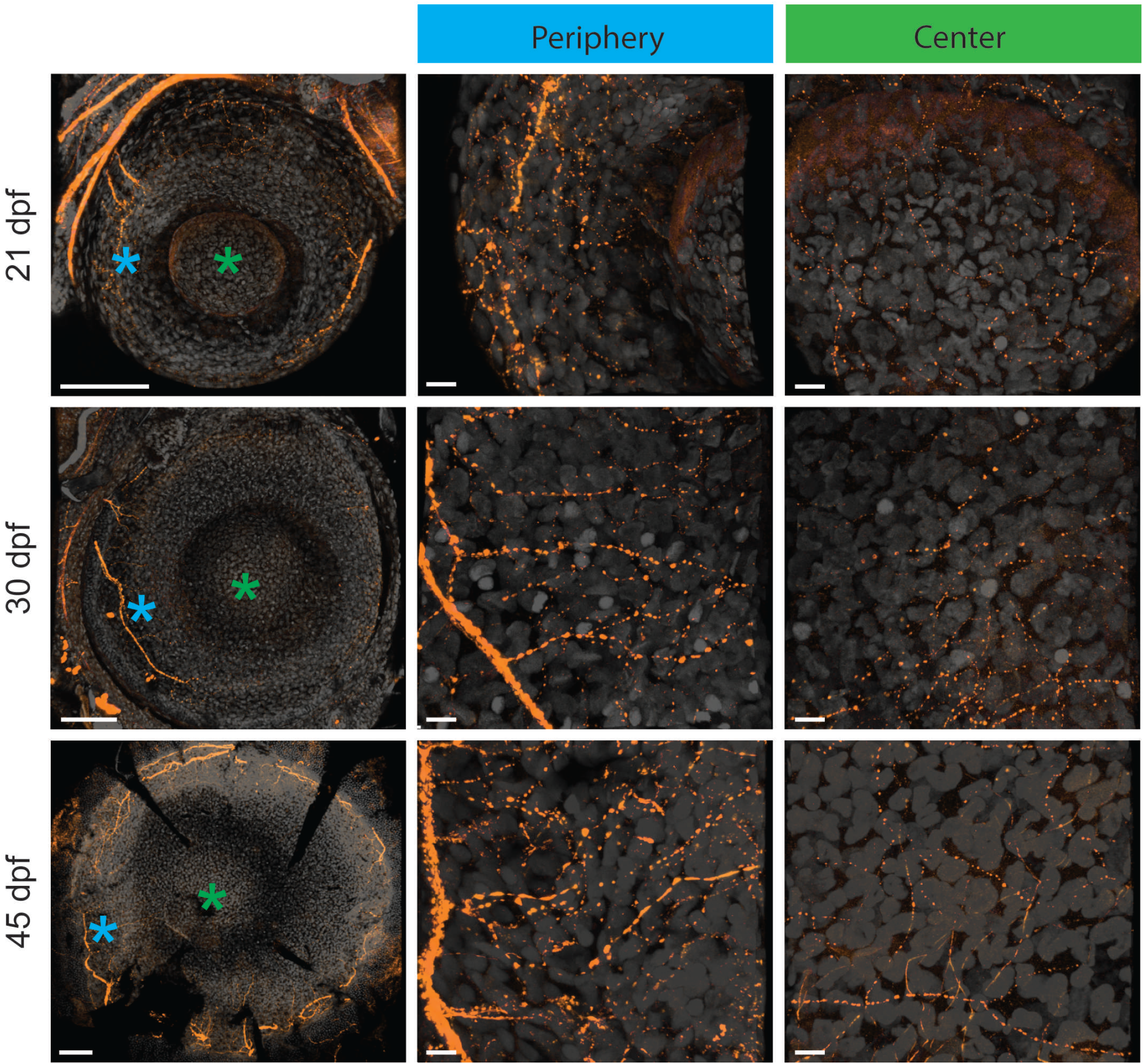
Corneal innervation during and after epithelial stratification at 21—45 dpf. Acetylated tubulin whole mount staining on maximum intensity projection images in central (area marked with green) and peripheral/limbal (area marked with blue) regions. In the periphery, the thickest branches extend to wider areas around the orbit, and branch out towards the central cornea. In central cornea, signal is re-gained. Scale bars: 100 µm in the overview images, 10 µm in the magnified images.

**Figure 8.**
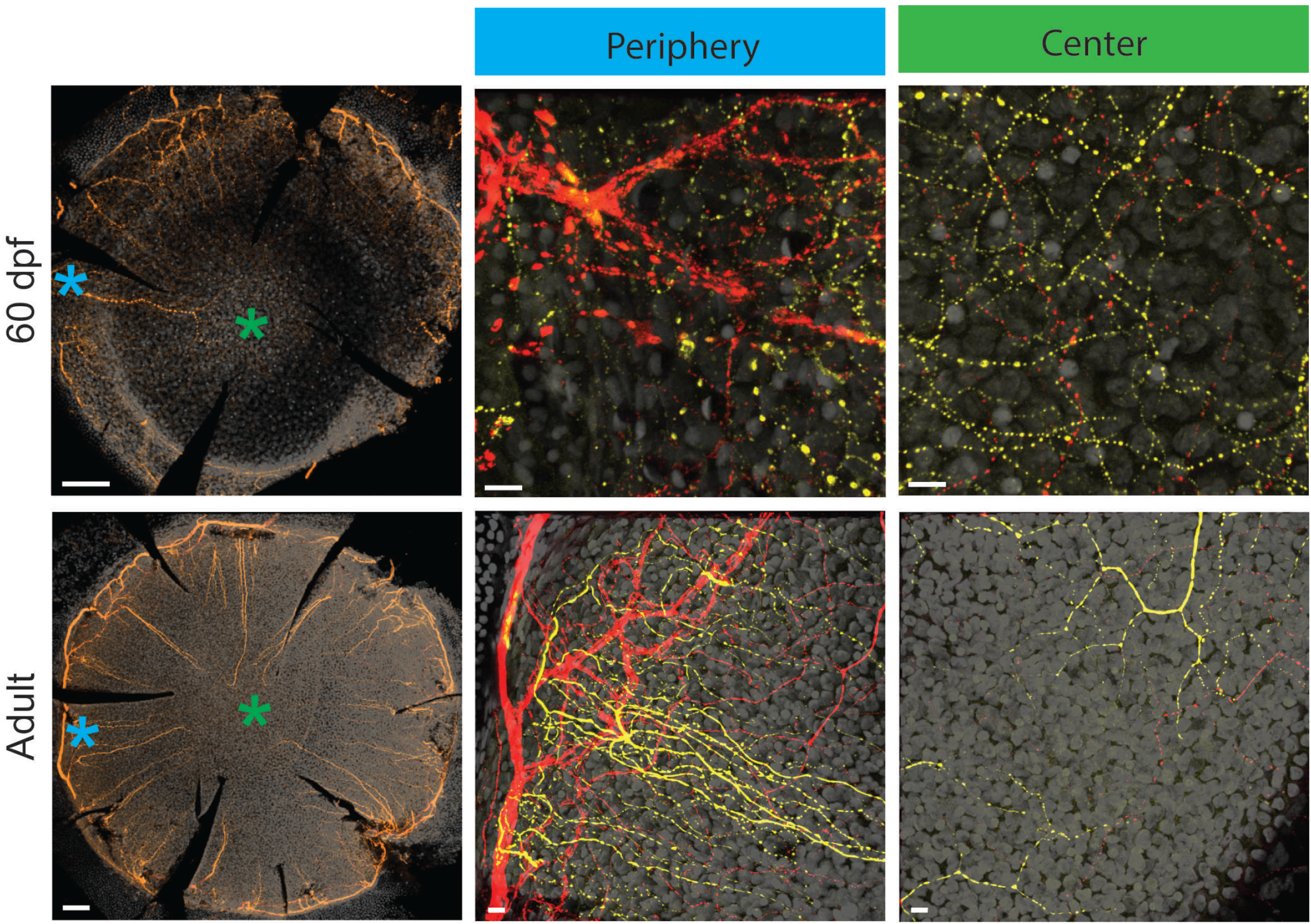
Corneal innervation in juvenile (60dpf) and adult fish. Acetylated tubulin whole mount staining on maximum intensity projection images in central (area marked with green) and peripheral/limbal (area marked with blue) regions. In these stages, the stromal (red) and epithelial (yellow) fibers can be distinguished. Scale bars: 100 µm in the overview images, 10 µm in the magnified images.

### Molecular maturation of corneal epithelium

The transcription factor Pax6 is expressed early on during eye development in presumptive ocular tissues of surface ectoderm and neuroectoderm origin (Nishina et al., 1999). In zebrafish, pax6 protein was reported on presumptive corneal epithelium from 48hpf on (Macdonald and Wilson, 1997). Here, we used RNAScope to detect *Pax6a* mRNA expression pattern, during the corneal maturation process. At 3dpf, the *Pax6a* expression was detected in the presumptive cornea (Fig. 9). As the signal was masked by *krtt1c19e* detection, we decided to present a staining of *Pax6a* alone (Fig. S2). The signal was, however, much lower than on the adjacent lens epithelium. From 7dpf to 21dpf, *Pax6a* expression was detected in most of corneal epithelial cells, at low level (Fig. 9). From 30dpf onwards, when the corneal epithelium is stratifiing, the *Pax6a* signal increased, and was detectable in all corneal epithelial cell layers.

**Figure 9.**
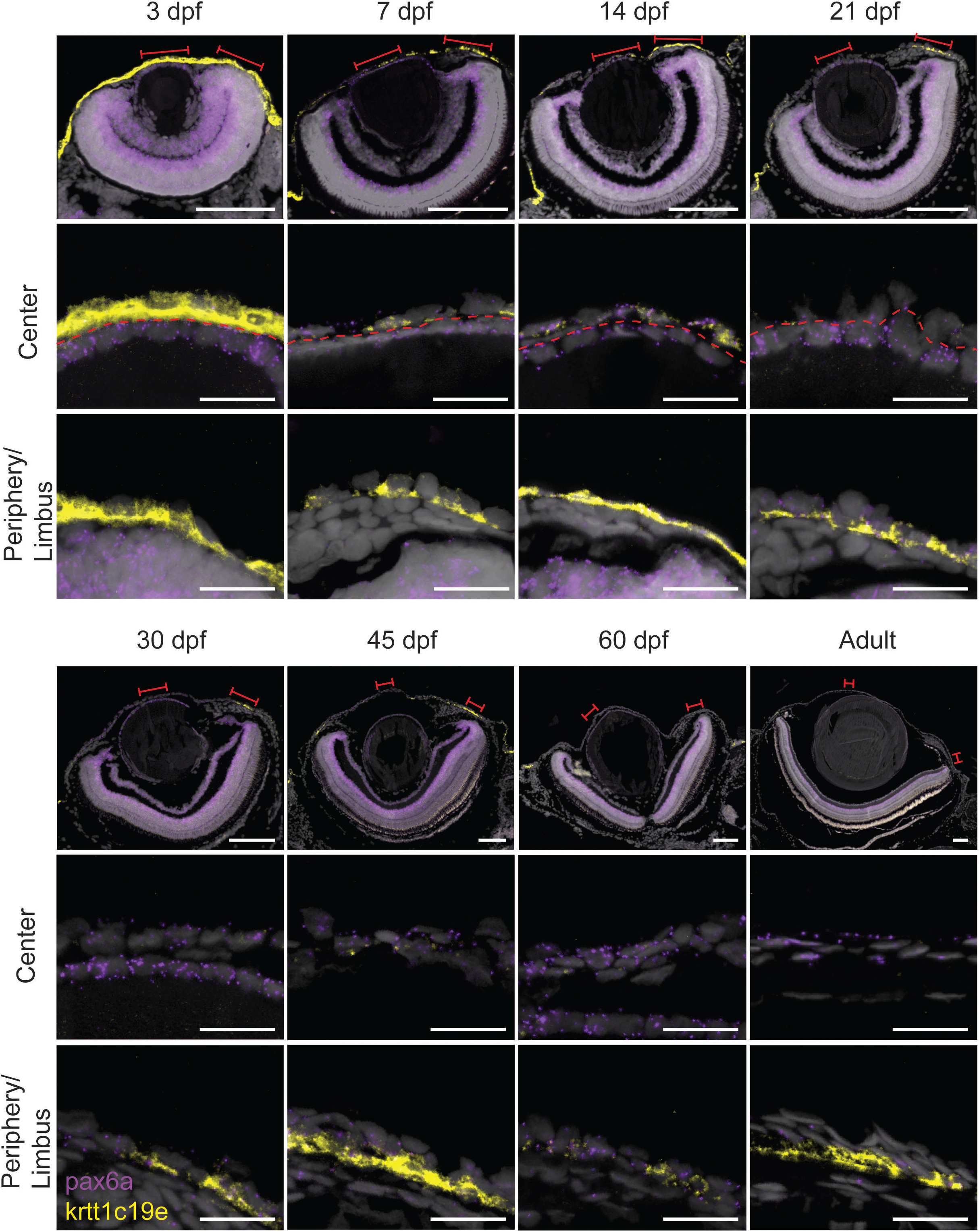
Expression of pax6a (red) and krtt1c19e (yellow) during corneal maturation. RNAScope in situ hybridization on 5-µm sections, red lines indicate areas in magnified images in center and periphery. Dashed line indicates the border between lens epithelium and cornea. Scale bars: 100 µm in the overview images, 20 µm in the magnified views.

In murine cornea, the level of maturation can be followed molecularly because of the switch from *keratin 14* (*krt14*) to *krt12* expression in central cornea (Kao, 2020). Therefore, in addition to *Pax6a* expression, we investigated the expression of a type I keratin, *krtt1c19e*, which has previously been reported to be present in the basal skin epithelium in zebrafish (Lee et al., 2014) (Fig 9). At 3dpf, we saw abundant expression throughout the surface of the embryo. From 7 dpf on, the central cornea showed only occasional *krtt1c19e* signal, whereas the peripheral regions retained its expression in the basal epithelial layer, even in the adult stage. In general, the peripheral cell population positive for krtt1c19e became smaller during maturation and growth. Interestingly, in fish 45 days old or younger, the posterior peripheral cornea seemed to have higher expression than the anterior peripheral region.

### Modulation of corneal epithelial markers during wound healing

The dynamics of *krtt1c19e* expression during corneal maturation seemed to reflect the acquisition of a specific cell fate in corneal periphery. We challenged this fate by wounding the corneal epithelium. Before the wound, *Pax6a* expression was found in close to all epithelial cells in the whole cornea, and *krtt1c19e* was predominantly found in the limbus, and few cells were positive in peripheral and central cornea (Fig. 10). We confirmed the results obtained previously (Ikkala et al., 2022), namely the loss of *Pax6a* expression in central cornea during the wound healing period. The *krtt1c19e* expression exhibited an interesting dynamics. 1hr post-wound (1hpw), we detected an increase of its expression spacially, as we found it in the periphery region, and in intensity in the limbus. Abruptly, at 6hpw, while the wound closed and the restratification was ongoing, *krtt1c19e* expression was totally lost and restarted slowly in limbal cells at 24hpw, reflecting going back to a physiological situation.

**Figure 10.**
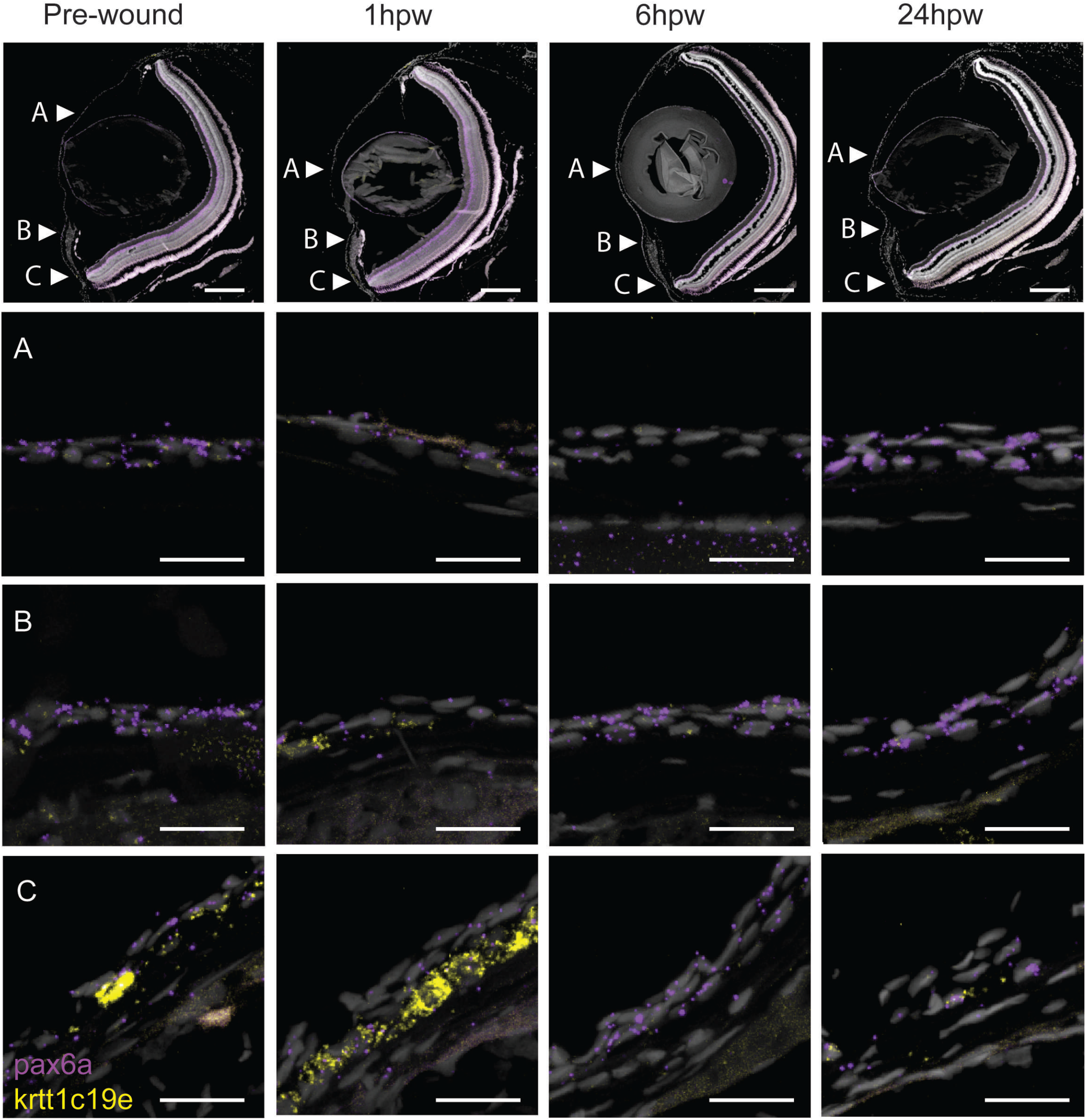
Gene expression changes in adult wound healing. RNAScope in situ hybridization on 5-µm sections showing the expression of *pax6a* (red) and *krtt1c19e* (yellow) before, and after 1, 6, or 24 hours, an epithelial abrasion on central cornea. Scale bars: 200 µm in the overview images, 20 µm in the magnified views.

## Discussion

Corneas found in camera-type eyes are structurally similar across animal kingdom, despite the various habitats colonized by animals bearing such eyes. Interestingly, the camera-type eye resulted from a convergent evolution in vertebrates and cephalopods (Serb and Eernisse, 2008). While the vertebrate cornea is a part of the eye, the cephalopod cornea is independent, and is often described as a specialized skin folding, which has an opening permitting the anterior chamber to be in contact with the sea water (Hanke and Kelber, 2019). Moreover, the corneal adaptation to various habitats is reflected in specific structural traits. For instance, our histology sections showed a thin stromal compartment when compared to murine cornea (Kalha et al., 2018). Strikingly, the stromal compartment in fish swimming in deep sea is thicker and more rigid than in mouse (Parravicini et al., 2012). Therefore, while the role of cornea in sight has been conserved, the structure was fine-tuned to adapt to various environmental conditions.

Among the most interesting traits of corneas, its maturation has always been fascinating. Numerous studies reported the late corneal maturation in mice. The *krt12* late expression, as marker of fully differentiated corneal epithelium, proves that cornea is fully mature at about 3 months of age (Tanifuji-Terai et al., 2006). In this process, *krt14* expression is confined to the limbal area, marking the less differentiated corneal epithelial cells. This dynamics is similar to the one of *krtt1c19e* that we described. Interestingly, at 45 and 60dpf, we still detected this keratin in epithelial cells, reflecting an incomplete maturation. Our data suggests that most of the epithelial maturation process happens between 14 and 45 dpf. Interestingly, our proliferation data showed that the limbal stem cells, which are label retaining cells, appeared between 21 and 30dpf. This time window corresponds as well to the apical cell size significant changes. Therefore, most of the changes within corneal epithelium take place fairly early on during zebrafish life.

Interestingly, mouse and zebrafish can both live for several years, about 3 and 5 years respectively. Therefore, we face a vast discrepancy in the timescale of corneal ontology, murine cornea being mature 140dpf and 45dpf for zebrafish cornea. Our current hypothesis is based on the time spent between birth and being weaned off. The postnatal and juvenile mouse is fed and protected, while the post hatching zebrafish larva, at about 2 dpf, relies on exploration and hunting, requiring a functioning sight (Privat and Sumbre, 2020).

Another element of cornea displaying a slow maturation is its innervation. In mouse, corneal innervation was reported to be fully mature by 4 months of age, which is a month after epithelium maturation completion (Bouheraoua et al., 2019). Our results showed a complete maturation of the corneal innervation in zebrafish between 2 and 6 months of age, which is later than the epithelium maturation. This feature of a delayed innervation maturation seems to be conserved. We could speculate that it might reflect the necessity of molecular and mechanical cues from the epithelium to complete the tissue innervation. The molecular cues could be some guiding factors that could be expressed after epithelium maturation, such as VEGF (Yu et al., 2008). The mechanical cues could be related to epithelial renewal, as in murine cornea, the swirl seen in epithelial nerves (Bouheraoua et al., 2019) mimicks the one seen in the epithelial renewal (Collinson et al., 2002).

Up to date, the innervation patterning mechanisms remain unknown. Strikingly, this pattern is not similar in all species. A recent report demonstrated that three main patterns exist in mammals, and the patterns are different in mouse, rabbit and pig (Marfurt et al., 2019). For instance, the rabbit corneal innervation pattern exhibits a horizontal patterning towards the inferonasal limbus, which is different to the epithelial renewal (Haddad, 2000). Our data showed a centripetal innervation pattern, similar to the epithelial renewal dynamics (Pan et al., 2013). However, we did not detect swirl, or vortex-like patterns in the central cornea. While we cannot exclude the possibility of missing a subpopulation of neurofilaments with our antibody, this option seems fairly unlikely as this antibody labels close to all corneal nerves in other species (Marfurt et al., 2019). Therefore, we can hypothesize that the guiding forces (molecular and/or mechanical), are comparable to the ones found in murine cornea, but might differ at the very centre of the tissue. Deciphering the cues responsible for corneal innervation mechanisms would bring invaluable new knowledge on corneal physiology, and help in understanding corneal pathologies, such as neurotrophic keratitis.

Taken together, our results present a much needed knowledge on zebrafish corneal biology. By deciphering the complete corneal ontogenic process, and maturation timeline, this report provides the fundamental insight required to use zebrafish cornea as a study model for upcoming research,

## Materials and methods

### Fish strains and maintenance

In this study we used the wild type AB fish line, acquired from the Zebrafish facility (HiLife, University of Helsinki). The fish were kept in 14:10 light-dark cycle in standard conditions in the Zebrafish unit. For each stage studied, the fish were randomly chosen from the stock tank. Adult stages refer to fish aged 4-9 months old. The experiments were done according to ESAVI/22167/2018 and ESAVI/1249/2022.

### Measurement of total fish length, eye diameter, and corneal growth

Fixed fish samples were placed on a plate, with millimeter paper underneath, and imaged. Images were opened in Fiji ImageJ 1.53. The scale was set using millimeter paper as reference, and total fish length and eye diameter (anterior-posterior) were measured with the line tool.

We used histological sections for measuring the growth of corneal epithelium. Stained sections from the middle of the eye were imaged with AxioImager M2, and the images were opened in Fiji ImageJ 1.53. The width of the cornea was measured, and normalized to anterior eye diameter (on the anterior-posterior axis).

### Scanning electron microscopy

Fish 3 – 45dpf (days post-fertilization) old were killed by prolonged anesthesia in 0.02% Tricaine solution (Sigma) on ice. 60dpf and adult fish were killed by anesthesia and decapitation. Fixing was carried out at +4°C in 2.5% glutaraldehyde (Grade 1, G7651, Sigma) in 0.1M sodium-phosphate buffer pH 7.4. After approximately 24 hours, samples were rinsed in 0.1M sodium-phosphate buffer pH 7.4. Eyes were dissected and stored in 0.1M sodium-phosphate buffer pH 7.4 at +4°C until further processing at the Electron Microscopy unit (University of Helsinki). Samples from fish 60dpf and older were treated with 2% osmium tetroxide (19130, Electron Microscopy Sciences), dehydrated in increasing ethanol series and by critical point drying. For smaller samples from younger fish, the final drying was done by letting the ethanol evaporate at RT. Finally, the samples were coated with platinum. Images were taken with FEI Quanta Field Emission Gun scanning electron microscope.

### Apical cell size measurement

We quantified the apical cell area on central and peripheral regions of the eye surface using the scanning electron microscopy images (2500X magnification) in Fiji ImageJ 1.53. The cell was manually outlined by the polygon tool, and the area was documented. Data were collected from three eyes per age group.

### EdU/BrdU-labeling

We labelled proliferating cells on two 24-hour time periods per group, using 0.2mM 5-ethynyl-2’’-deoxyuridine, EdU (900584, Sigma), in time point 1, and 0.2mM 5-bromo-2’-deoxyuridine, BrdU (B5002, Sigma) in time point 2. The labeling reagent was diluted in E3 buffer or system-water, depending on the age of fish, from 10mM (EdU) or 8mM (BrdU) stock prepared in double-distilled water and stored at -20 °C.

### Corneal abrasion

The fish were anesthetized in 0.02% Tricaine and placed into an incision on a moist sponge, head protruding from the sponge surface. The eye surface was abraded with an ophthalmic burr (Algerbrush II, BR2-5, Alger Equipment Company) with a 0.5-millimeter tip, by pressing the rotating burr tip gently onto the eye and moving it with circular motion. Then the fish was placed back to the tank water.

### Whole mount staining

Fish were euthanized as above, and tissue fixed for 20 minutes on ice in 4% PFA (15713, Electron Microscopy Sciences) in PBS. Then, samples were rinsed with PBS and stored in 100% methanol at -20 °C. Upon staining, samples went through rehydration steps in decreasing methanol series (75/50/25% MetOH/PBS, 5 min each) at room temperature, and PBS rinsing. The eyes of adult fish were enucleated at this point. Samples were permeabilized in 0.5% Triton-X-100 (807423, MP Biomedicals) in PBS for one hour at roo temperature, and blocked (10% goat serum (16210064, Life Technologies) + 0.5% BSA (A2153, Sigma), in 0.1% Triton/PBS) for 3—6 hours at room temperature, and incubated in primary antibody overnight at +4 °C (1:200 in blocking solution). We used the primary antibody mouse-anti-acetylated tubulin (T7451, Sigma). Samples were then washed three times for one hour at room temperature in 0.1% Triton/PBS, blocked for 1—2 hours at room temperature, and incubated in secondary antibody (goat-anti-mouse IgG, Alexa 568, A11057, Life Technologies) diluted 1:200 in blocking solution over night at 4 °C. Finally, samples were washed as above, rinsed in PBS, counterstained in Hoechst (H3570, Invitrogen) 1:2000 in PBS for 30 minutes at room temperature, and mounted. For young stages, we mounted the tissue in 1% low-melting-point agarose (R0801, Life Technologies) on imaging dish. For fish ≥45dpf old, corneas were dissected and flat-mounted in 80% glycerol on microscopy slides.

### Stainings on formalin-fixed, paraffin-embedded sections

Fish were euthanized as above, and fixed overnight in 4% PFA/PBS at +4 °C. Samples were then embedded in Histogel (HG-4000, Thermo Fisher), and subjected to automated dehydration, xylene incubation, and paraffin embedding using the ASP 200 tissue processor machine (Leica Biosystems). Samples were sectioned with 5-*μ*m thickness on the coronal plane, dried overnight at +37°C, and baked briefly on a hotplate. Upon staining, sections were rehydrated in decreasing ethanol series, permeabilized in 0.5% Triton/PBS for 10 minutes, and subjected to heat-induced antigen retrieval in automated retriever machine (62700, Electron Microscopy Sciences) in 10 mM sodium-citrate buffer, pH 6.0. The sections were then washed in PBS for 10 minutes, blocked for one hour (10% goat serum in 0.1% Triton/PBS), and incubated with mouse-anti-BrdU (RPN202, Cytiva) 1:100 in blocking solution overnight at room temperature. On the following day, the sections were washed in 0.1% Triton/PBS for 15 minutes, incubated in secondary antibody (goat-anti-mouse Alexa 568) 1:200 in blocking solution for 2 hours at RT, washed in 0.1% Triton/PBS and PBS, both for 15 minutes. Sections were incubated in 3% BSA/PBS for 10 minutes. EdU tracing was done with the kit reaction components according to kit instructions (C10337, Invitrogen). Sections were then incubated again in 3% BSA/PBS for 10 minutes, washed in PBS for 10 minutes, incubated in Hoechst 1:2000/PBS for 20 minutes, washed in PBS for 10 minutes, and mounted in Fluoromount-G (0100-01, Southern Biotech).

### RNAScope *in situ* hybridization

We detected the expression of pax6a and krtt1c19e with the RNAScope Fluorescent V2 kit (323110, BioTechne), using probes Dr-pax6a (532481) and Dr-krtt1c19e (1117231-C2), and Opal dyes 520 (1:1000 dilution) and 620 (1:1500 dilution) (FP1487001KT, FP1495001KT, Akoya Biosciences). We performed the staining on 5-micrometer paraffin-embedded, formalin-fixed sections, using manufacturer’s protocol with an additional baking of sections at 60°C for 30 minutes after deparaffinization. We performed a 17-minute target retrieval, followed by a 20-minute protease treatment.

### Light microscopy and image processing

The bright-field images on Figure 1 were obtained with Zeiss Axio Imager M2. The white balance and contrast were improved on whole images in Adobe Photoshop (Version 22.5.4). The images were re-scaled, cropped, and placed into figure panel in Adobe Illustrator (Version 25.4.1).

The fluorescently labeled samples were imaged with Leica SP8 inverted confocal microscope (Leica Microsystems, Wetzlar, Germany), by acquiring image stacks, and tile scans when necessary, with HC PL APO 10x/0.40 CS2, HC PL APO 20x/0.75 CS2, or HC PL APO 63x/1.20 CS2 objective. The excitation/emission detection wavelengths were: 405/430—480 (Hoechst), 488/500—550 (Alexa 488, Opal 520), 561/570—650 (Alexa 568, Opal 620).

Signal intensity and channel colors were adjusted in Imaris software (Bitplane), and snapshots used for image panels. The epithelial and stromal compartments in Figure 8 were defined by the basal epithelial cell layer, which can be recognized by the even distribution and shape of the nuclei.

### Statistical analysis

Numerical data within a figure was tested for normal distribution. We used ordinary one-way ANOVA for normally distributed data, with Sidak’s multiple comparisons test. For data not normally distributed, or with small n-number, we used Kruskal-Wallis test with Dunn’s multiple comparison test. All statistical testing was done with Graphpad Prism (8.3.0).

## Captions

**Figure S1**. Central cornea appearance in 21dpf fish. The dashed line points to the area presented in higher magnification. Scale bars: 20 µm.

**Figure S2**. The *pax6a* signal on 3dpf cornea. A. Overview of a coronal section of the eye. B. Central region of the eye. Dashed line separates the lens and the cornea C. Peripheral region of the eye. D. Adjacent tissue. Le=lens epithelium, Ce=corneal epithelium. Scale bars: 50 µm in A, 10 µm in B-D.

