## Supplementary figures and images for "Zebrafish cornea formation and homeostasis reflect robustness in camera-type eye biology, regardless of its environment"

### Supplemental Figures

#### Figure S1

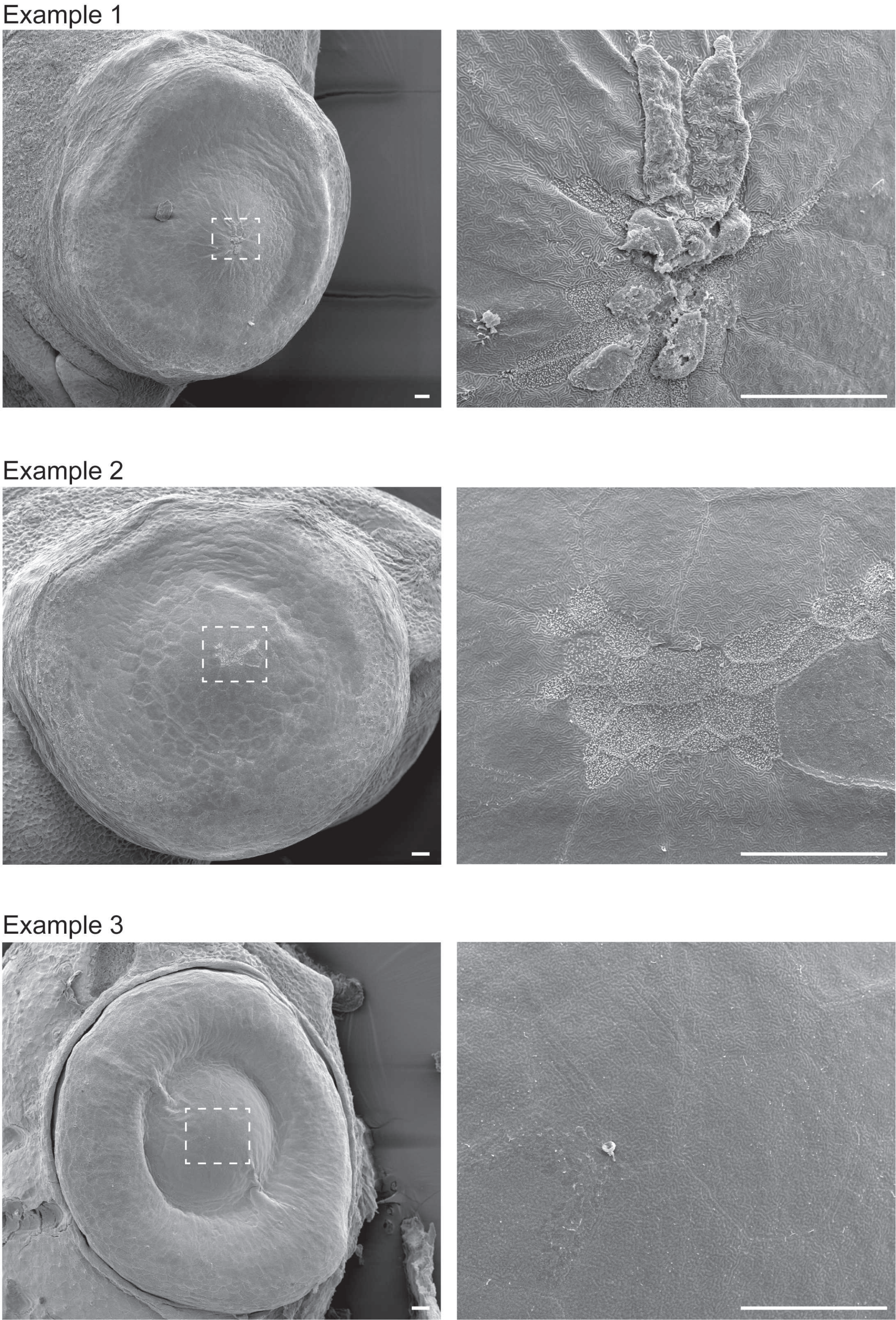

Figure S2

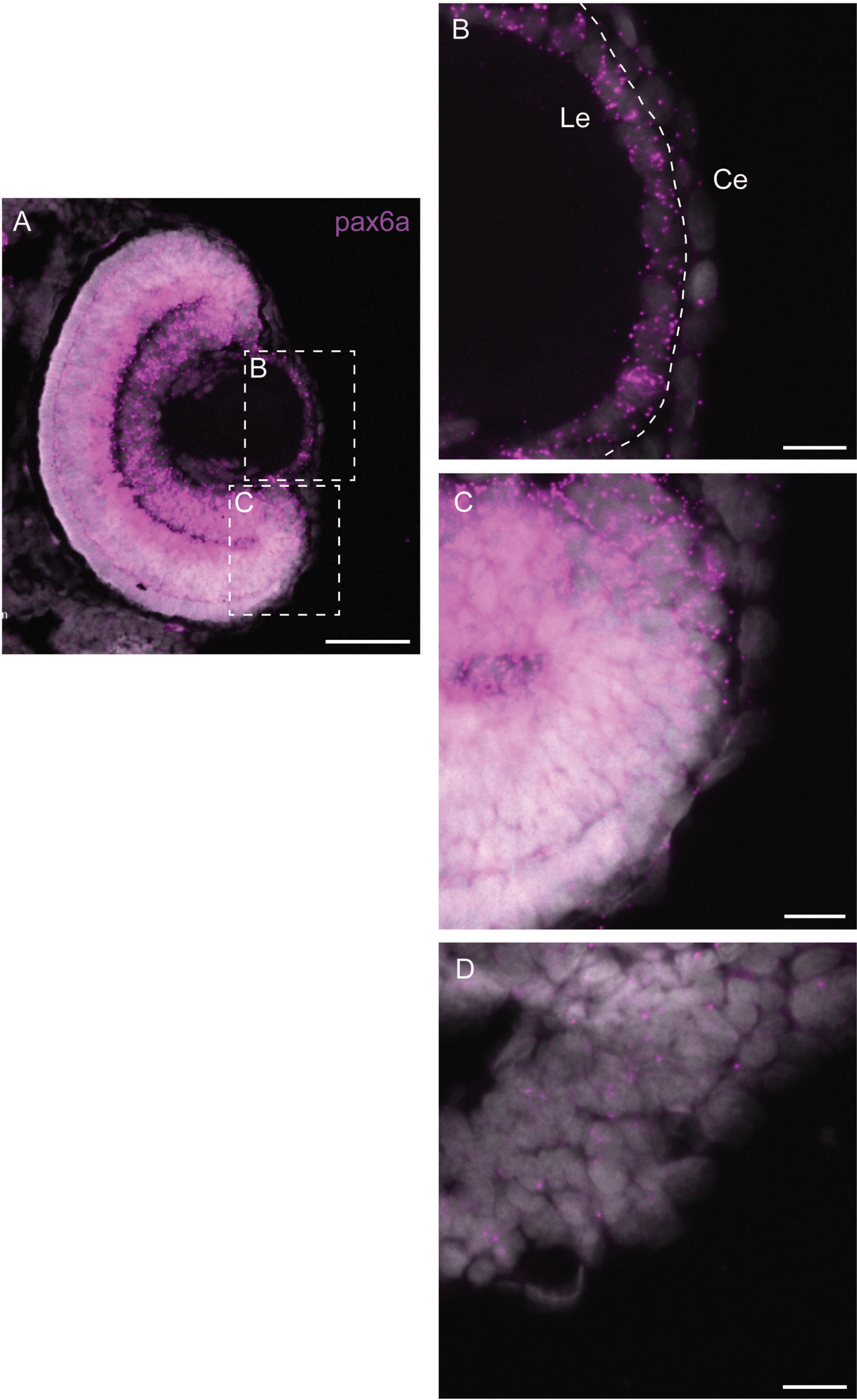
